# Elucidation of the viral disassembly switch of tobacco mosaic virus

**DOI:** 10.1101/569178

**Authors:** Felix Weis, Maximilian Beckers, Iris von der Hocht, Carsten Sachse

## Abstract

Stable capsid structures of viruses protect viral RNA while they also require controlled disassembly for releasing the viral genome in the host cell. A detailed understanding of viral disassembly processes and the involved structural switches is still lacking. Biochemically, this process has been extensively studied using the tobacco mosaic virus model system and carboxylate interactions have been proposed to play a critical part in this process. Here, we present two cryo-EM structures of the helical TMV assembly at 2.1 and 2.0 Å resolution in conditions of high Ca^2+^ concentration at low pH and in water. Based on our atomic models, we identified the conformational details of the disassembly switch mechanism: in high Ca^2+^/acidic pH environment the virion is stabilized between neighboring subunits through carboxyl groups E95 and E97 in close proximity to a Ca^2+^ binding site. Upon increase in pH and lower Ca^2+^ levels, mutual repulsion of the E95/E97 pair and Ca^2+^ removal destabilize the network of interactions at lower radius and release the switch of virus disassembly. Our TMV structures revealed the conformational details for one of the reference systems of viral assembly/disassembly and provide the mechanistic explanation of a plethora of experimental results that were acquired over decades.

**Significance Statement:** Tobacco mosaic virus presents the text-book example of virus structure and RNA release from a viral capsid through disassembly. Despite the wealth of structural and biochemical data on the assembly/disassembly properties generated from more than 80 years of research, the atomic-resolution structural details of the proposed conformational changes have not been resolved to date. The here determined high-resolution cryo-EM structures reveal the conformational details of the molecular disassembly switch. When the virus enters the cell, carboxylate repulsion and loss of calcium-ion coordination destabilize the switch region and can trigger RNA release through virus disassembly. The two determined structural states resolve a long-standing question on environment-driven virus disassembly switches.

## Introduction

The capsid of RNA viruses provides a container that ensures the protection of the viral genome from degradation in the extracellular environment. The shell of tobacco mosaic virus (TMV) is particularly thermostable in ambient temperatures and resistant to degradation across a wide range around neutral pH [1]. The TMV capsid is primarily composed of a 17kDa coat protein (CP) that is organized in a helical assembly thereby tightly enclosing the viral RNA of the genome [2]. The mature virion forms a rigid rod of approximately 300 nm length resembling a hollow cylinder with an inner and outer diameter of 4 and 18 nm, respectively. A single helical turn is built from 16 1/3 CP subunits making up the pitch of 23 Å. These straight helical structures were subject to pioneering studies in our understanding of biological matter [3][4] and due to high stability and easy availability. TMV is still a widely used model system both in virology as well as structural research [5][6][7][8]. Increasingly, the virus is becoming of interest to future hybrid biotechnological applications at the interface with material science [9].

In order to propagate the virus during the life cycle, the capsid protein requires controlled opening inside the host cell in order to release the RNA and initiate translation for replication. This way of viral entry requires a molecular mechanism that is capable of sensing the differences between the extracellular and the intracellular medium, followed by destabilization of the assembled virion. A series of plant viruses use Ca^2+^ and pH triggered disassembly [10] and thereby exploit the lower Ca^2+^ and proton concentrations inside the plant cell compared with the extracellular environment. Many of these findings were observed in early studies of TMV [10][2][3]. TMV is capable of entering the cell through mechanical lesions that transiently open the outer membrane [13].

The disassembly behavior of TMV has been biochemically studied in remarkable detail. Based on titration experiments, it was reported that TMV contains groups that titrate with pK_a_ values between 7 and 8 [12], leading to the hypothesis that a carboxylate cluster binds protons with high affinity. These putative residues termed “Caspar-Carboxylates” [12][14] are thought to drive virus disassembly through mutual repulsion upon entering the intracellular environment. In addition, it has also been demonstrated that TMV binds Ca^2+^ via these residues [15][16]. Mutational studies have identified the critical residues [12][18] involved in the disassembly process: E50 and D77 at medium radius of the cylinder cross section have been hypothesized to be involved in axial carboxylate interactions, whereas E95, E97 and E106 at lower radius mediate lateral carboxylate and possible Ca^2+^ interactions.

Despite the wealth of biochemical and mutational studies, our structural understanding of the conformational switch sensing the environmental changes is still incomplete. Although TMV was subject to a plethora of structural studies [5][19][20] [21], resolution of the helical rod was limited to 2.9Å when determined by early X-ray fiber diffraction [2] or later to 3.3 Å by electron cryo-microscopy (cryo-EM) studies [5][22]. At these resolutions, the density was sufficiently clear for the assignment of the architecture of the CP and annotation of bulky sidechains, whereas further details regarding the conformation of the implicated side chains remained undetermined. The proposed Caspar carboxylate residues E95, E97, E106 and the calcium binding site are found in a more flexible part of the protein at lower radius with high B-factors, as it is expected for such a metastable switch. In the absence of RNA within the disk assembly of TMV at higher resolution the respective residues were not detectable and assumed to be disordered [23]. Generally, negatively charged amino acid residues suffer from faster radiation damage when imaged by cryo-EM [24][5], which makes them more difficult to model. Therefore, the precise structural details of this intricate viral disassembly switch remain to be elucidated.

In order to address this outstanding fundamental question regarding viral disassembly, we used cryo-EM including latest developments of high-resolution imaging, data processing and density interpretation methods and determined two ∼2 Å resolution TMV structures. The densities reveal that the metastable switch is based on a Ca^2+^ sensitive network of carboxylate and iminocarboxylate residues at lower radius, which become destabilized by Ca^2+^ release at higher pHs. The cryo-EM structures captured TMV at different conformational states of this network and thereby directly reveal the transitional mechanics of the switch driving viral disassembly.

## Results

### Cryo-EM structures of TMV at 2.0 and 2.1 Å resolution

To determine the cryo-EM structures of TMV and elucidate different structural states of the viral assembly, we prepared TMV in two different conditions, first in the presence of 20 mM CaCl_2_ at pH 5.2 (referred to as Ca^2+^/acidic pH) and second in water in the absence of any cations. Both samples were plunge-frozen and imaged using a 300kV electron microscope including a direct electron detector (Figure 1a). With the collected micrographs, we determined the 2.1 and 2.0 Å resolution TMV structures in Ca^2+^/acidic pH and water, respectively (Supplementary Figure 1a). Both maps show local resolutions up to 1.8 Å for the CP core and ∼5Å for the disordered C-terminal tail (Figure 1b, Supplementary Figure 2). Map details agree with the expected high-resolution features such as defined carbonyl oxygens of the protein backbone. We identified a total of 101 water molecules for TMV in water, 83 water molecules under Ca^2+^/acidic pH, 4 Mg^2+^-ions bound to RNA as well as well-defined side-chain conformers per CP in both conditions (Figure 1c, d, e). A critical Ca^2+^ could be located in the Ca^2+^/acidic pH structure whereas no Ca^2+^ binding was observed at the proposed Ca^2+^ site at the RNA [2][22] in both maps (Supplementary Figure 3a). In order to verify the principal biological activity of the imaged virus structures, we demonstrated infectivity of the used virus batch in tobacco plants, which showed typical symptoms like stunted growth and necrotic lesions 35 days post-infection (Supplementary Figure 4).

**Figure 1.**
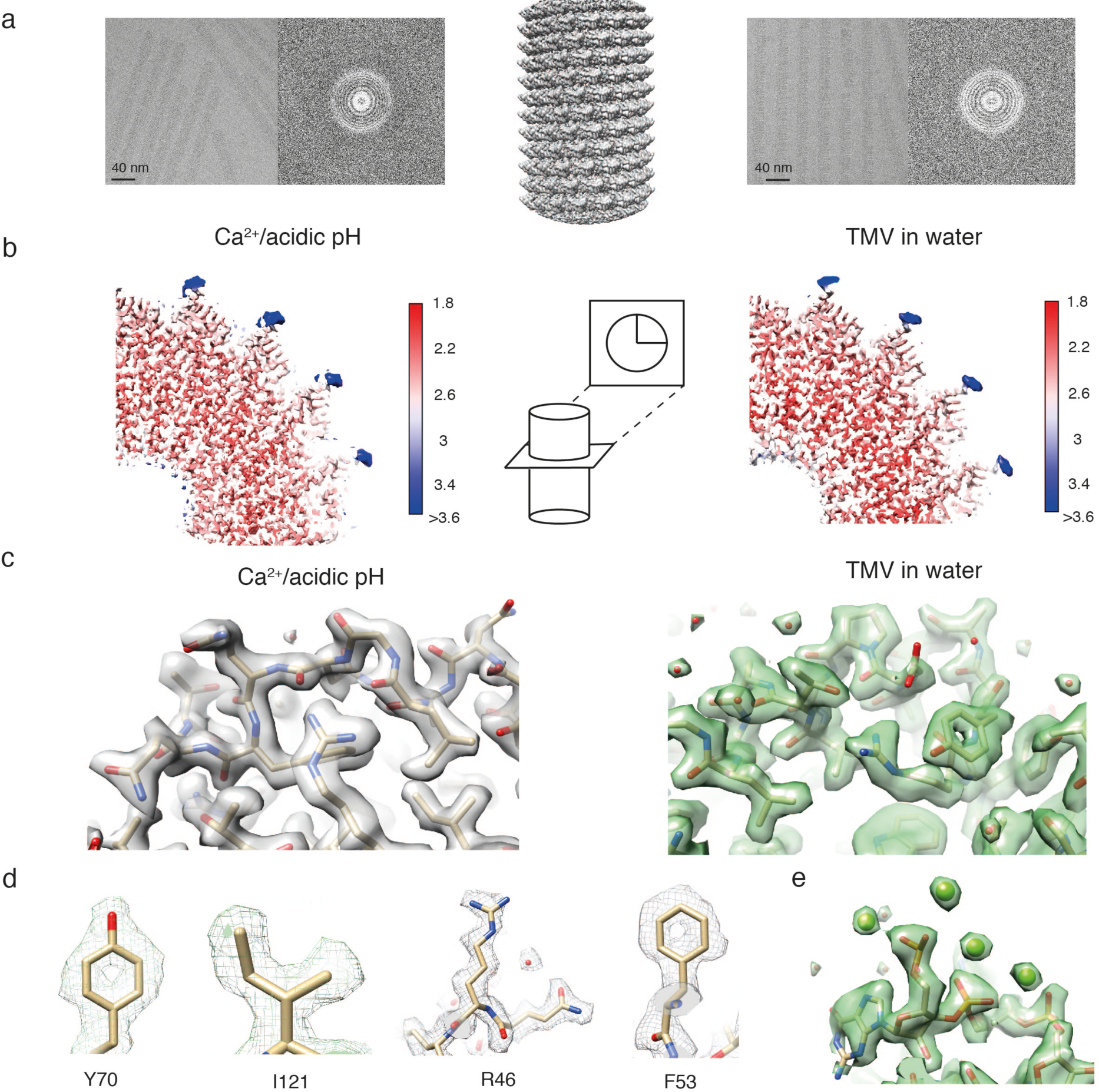
High-resolution cryo-EM structures of tobacco mosaic virus (TMV) in conditions of Ca^2+^/acidic pH (grey, left) and water (green, right). (a) Characteristic micrographs side by side with corresponding power spectra for both data sets, respectively (left and right). (b) Local resolutions mapped on the respective 3D reconstructions. The center of the coat protein is resolved up to 1.8 Å with some features of approx. 5 Å resolution at the C-terminus. (c) Both maps show well-resolved protein features including water molecules (left, right). (d) Side-chain features of Y70, I121, R46 and F53 residues at Ca^2+^/acidic pH. (e) Snapshot of RNA density with Mg^2+^ ions in water conditions.

### Ca^2+^/acidic pH and water structures at lower radius

Using our recently developed statistical framework for the annotation of molecular density [25], we were able to consistently analyze the CP density from Ca^2+^/acidic pH and water samples (Figure 2a). Although the two determined TMV maps are very similar for most of the CP, the lower radius region differs significantly (Figure 2a **right**). Comparison of this lower radius density with recently determined EM densities EMD2842 [5] (Supplementary Figure 2b) revealed that previous studies only poorly resolved this part of the protein as the density values were much weaker than other parts of the protein. Detailed comparison of the lower radius densities revealed that the protein backbone follows an alternative path in the Ca^2+^/acidic pH and water structure (Figure 2b **top**). In the determined Ca^2+^/acidic pH structure, we built an atomic model matching the density (Figure 2b **left**). In the water structure at lower radius, however, we identified 3 co-existing models in the residue range 97 – 100 that describe the density, e.g. the density of the water structure is consistent with multiple conformations of E97 (Figure 2b **right**). Closer inspection of the lower radius interface between neighboring subunits reveals additional differences: in the Ca^2+^/acidic pH structure, density of a bound ion is present and consistent with coordination by E106, N101, N98 and a backbone carbonyl oxygen. In the water structure, however, this complete coordination is missing and asparagines N101 and N98 are facing away from the central density. Therefore, we conclude that under Ca^2+^/acidic pH conditions this subunit interface is stabilized by a Ca^2+^ ion, whereas in water it is replaced by H_2_O molecules.

**Figure 2.**
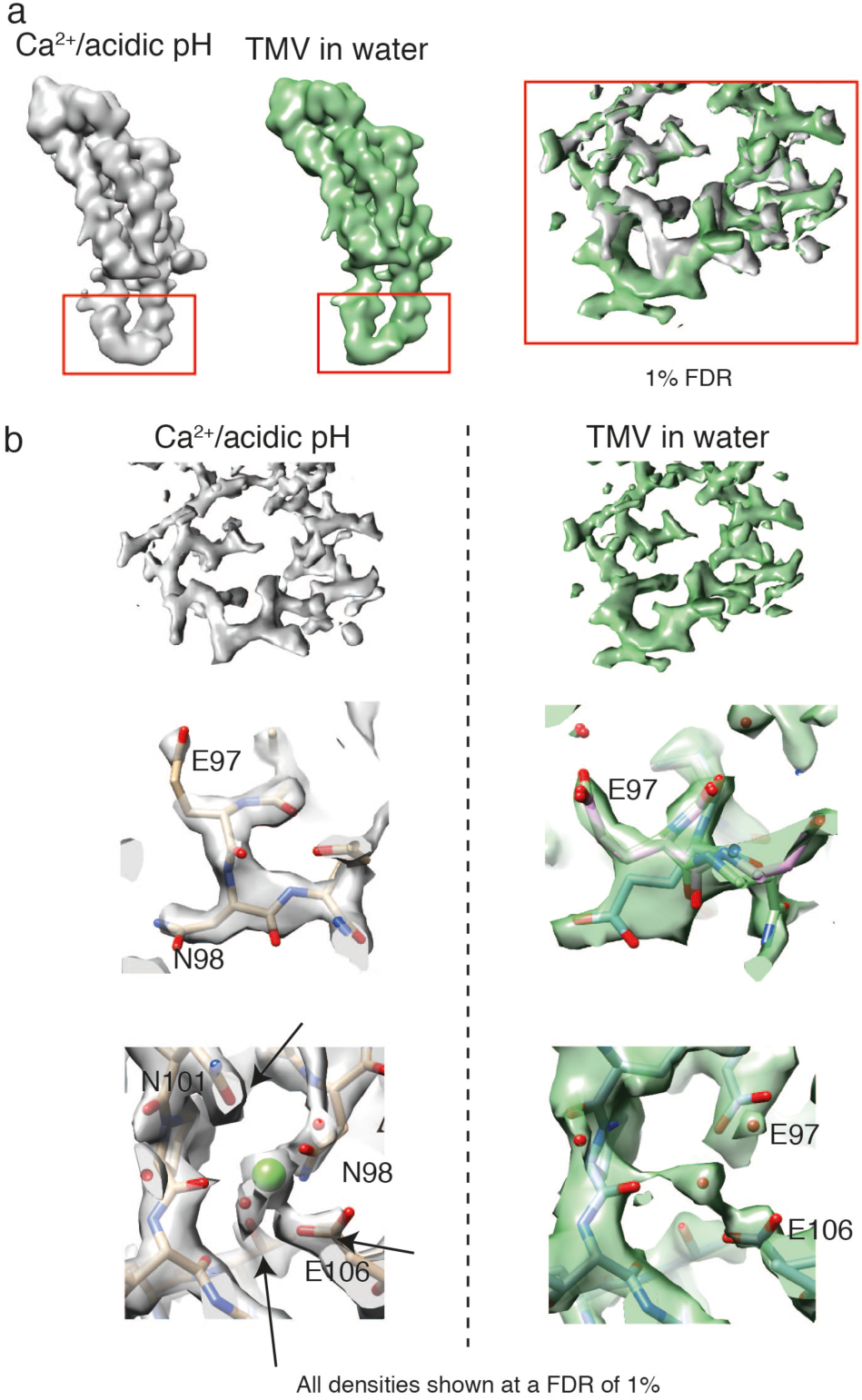
Confidence maps thresholded at 1 % FDR from cryo-EM maps of Ca^2+^/acidic pH and water structure. (a) Low-pass filtered monomer densities of Ca^2+^/acidic pH (grey) and water structure (green) with differences at lower radius (left). Zoomed inset (right) with maps displayed at a false discovery rate (FDR) threshold of 1% showing significant differences. (b) Detailed map comparison at lower-radius region of Ca^2+^/acidic pH (grey, top row) with the water condition (green, bottom row). A total of 3 atomic models (model 1: cyan, model 2: pink, model 3: green) describe the density of TMV in water whereas the Ca^2+^/acidic pH density could be modelled with a single atomic model (center). Located Ca^2+^ ion in the Ca^2+^/acidic pH density with different conformations in the water structure (bottom).

### Interactions involved in the metastable switch

In order to analyze the detailed interactions within the two structures in more detail, we refined the atomic coordinates by a common real-space optimization approach [26]. Comparison of the refined atomic models revealed the stabilization of an additional α-helical segment in the Ca^2+^/acidic pH model between E97 and A100 (Figure 3a). Plots of refined atomic B-factors also show a decrease from 42 to 30 Å^2^ in the lower radius region for the Ca^2+^/acidic pH condition supporting the notion that Ca^2+^ stabilizes the assembly structure in comparison with water (Supplementary Figure 5). Next, we more closely examined the carboxylate residues previously identified to be critical in the disassembly process (E50, D77, E95, E97 and E106). First at medium radius, E50 and D77 contribute to tight axial carboxylate contacts at a distance of 2.6 Å and showed no differences between the determined structures. Second at lower radius, we find that glutamates E97 and E95 make up tight inter-subunit interactions in the Ca^2+^/acidic pH structure with a distance between the carboxylate groups of 3.5 Å (Figure 3b **left, top**). Although E106 is not found in contact with other carboxylates, E106 is involved in the coordination of Ca^2+^. The respective Ca^2+^ site is coordinated by E106 and N98 from one CP monomer and N101 as well the backbone carbonyl oxygen of P102 from the neighboring monomer (Figure 3b **left, bottom**). These close inter-subunit interactions between neighboring CPs add to the stability of the helical assembly. The water structure, however, is lacking the close inter-subunit carboxylate contacts as E97 shows two different conformations, one facing towards E106 (model 1) and the other towards E95 (model 2) with significantly longer distances of 4.7 and 5.3 Å between carboxylates, respectively (Figure 3b **right, top**). Residues N101 and N98 also assume different conformations in the water structure (Figure 3b **right, bottom**), whereas they participate in the coordination of Ca^2+^ in the Ca^2+^/acidic pH condition. We conclude that due to the loss of Ca^2+^ coordination as well as the buildup of carboxylate repulsion at higher pH, the residue network in proximity of E106, N101 and N98 becomes destabilized and changes conformations. (Figure 4).

**Figure 3.**
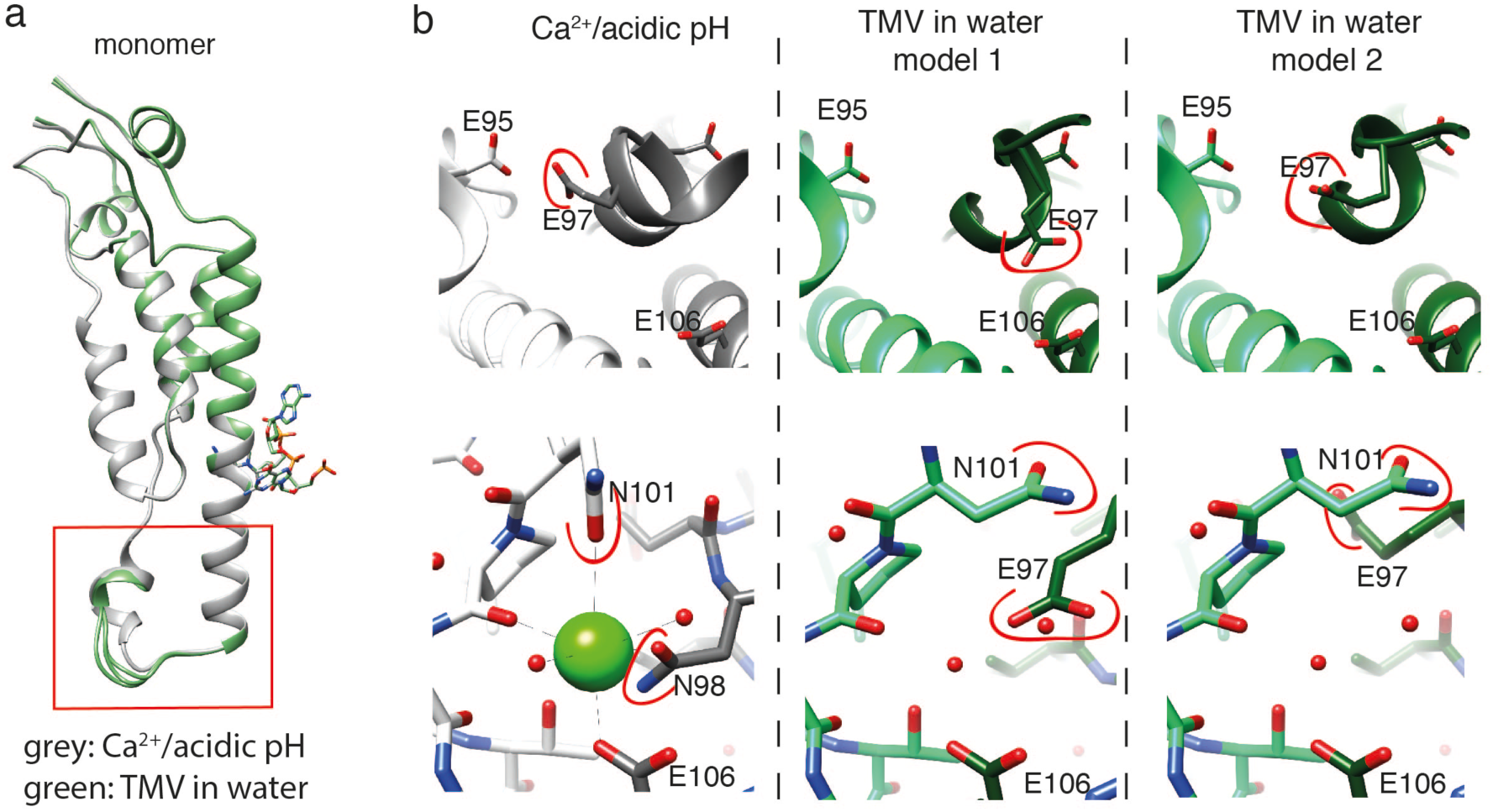
Model comparison of Ca^2+^/acidic pH and water structural states. (a) Superpostion of the monomer structures with the lower radius region highlighted in the red box. The Ca^2+^/acidic pH state (grey) shows an additional α-helical segment when compared with the water models (green). (b) Comparison of the E95-E97 interaction (top row) and of the Ca^2+^ binding site in the Ca^2+^/acidic pH model (bottom row). Close proximity of E95 and E97 in the Ca^2+^/acidic pH state whereas in water E97 flips towards E106 and is rather flexible. No Ca^2+^ ion and corresponding coordination is evident in the water model. Residues that change conformation are marked in red. Adjacent subunit models are displayed in lighter shade of the main color. Model 3 is not shown due to high structural similarity with model 2 in the displayed region.

**Figure 4.**
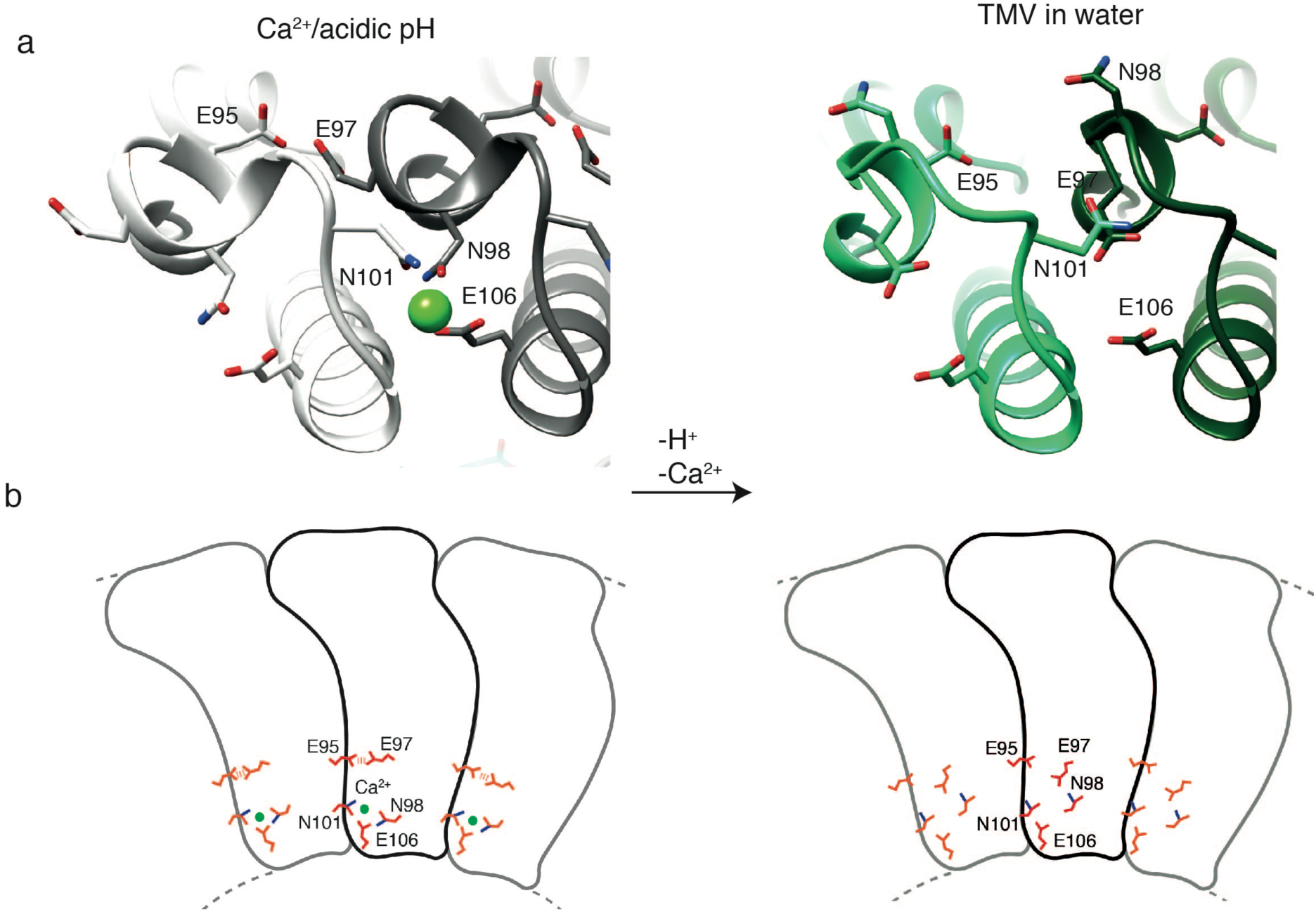
Towards a disassembly mechanism based on the Ca^2+^/acidic pH and water states. (a) The close proximity of the E95-E97 interaction and tight coordination of Ca^2+^ in the Ca^2+^/acidic pH state site suggests a mechanism in which, upon pH change and Ca^2+^ removal, repulsive forces between charged carboxylates destabilize the network of interactions at lower radius releasing the switch for viral disassembly. (b) Schematic presentation of coat protein with neighboring subunits including main residues of TMV in the Ca^2+/^acidic pH state (left) and in water (right) responsible for the metastable disassembly switch.

## Discussion

In this study, we demonstrated that structure determination by cryo-EM is capable of resolving helical specimens to resolutions up to 2.0 Å. Statistical analysis combined with local resolution assessment [25] allowed density annotation of the complete CP including disordered residues and post-translational modifications. Furthermore, for the first time, we could interpret the critical residue conformations at the lower radius region that so far evaded visualization at previous 3.3 Å resolution structures due to higher flexibility and radiation damage. In the 1960s, critical glutamate residues at lower radius were proposed to drive virus disassembly [2] and since then have often been referred to as “Caspar carboxylates”. Based on two high-resolution cryo-EM structures in the presence and absence of Ca^2+^, we propose a detailed structural mechanism of virus disassembly. In the presence of Ca^2+^ at acidic pH, tight inter-subunit interactions via Ca^2+^ coordination and carboxyl-carboxyl(ate) interactions between E95 and E97 stabilize the assembly. In these conditions, carboxyl residues are able to bind protons and neutralize their negative charges, which weakens their repulsive force leading to close contact between E95 and E97 (Figure 4). Upon entering the cell, the pH rises and carboxyl groups are deprotonated leading to repulsive forces between them. Putative movement of E97 away from E95 and correlated motion of N98 and N101 destabilize the ion binding site and further promote Ca^2+^ removal in a low Ca^2+^ environment. These concerted conformational rearrangements loosen the stabilizing inter-subunit interactions and ultimately release the switch to disassemble the virus (**Supplementary Movie**). The involvement of additional residues such as N101 and N98 in the coordination of the Ca^2+^ binding site suggests a more intricate conformational network responsible for rearrangements beyond the previously postulated carboxylate repulsion driving disassembly, which is corroborated by results from a series of mutation experiments [17]. It should be noted that due to the convoluted nature of conformational changes at the lower radius regions, it is not possible to assign a temporal order to the conformational changes from the two observed structural states of TMV assembly.

Previous studies proposed a critical Ca^2+^ site that interacts with the RNA backbone [2][22]. Such a site could not be located in our two structures (Supplementary Figure 3a). In fact, the conformations around the RNA in both our structures resemble what has been referred to as the low Ca^2+^ state [22], with D116, R92 and R90 involved in RNA binding [21] and the direct interaction of R92 and D116. According to our structures, this previously proposed second Ca^2+^ site [2][22] and the different conformation at the RNA may not be required for TMV stabilization. In addition, residue D109 thought to be important for disassembly was not found to assume different conformations in the two structures and did not form interactions with one of the before mentioned residues (Supplementary Figure 3b). To what extent the noted structural differences reflect the different preparation conditions or experimental uncertainties of the previous atomic models is not easy to resolve. Our 2.0 and 2.1 Å resolution cryo-EM maps are sufficiently clear to locate all the mentioned side chains with high confidence (Figure 3). In order to confirm that our batch of TMV presents a biologically active virus, we also showed experimentally that our sample is capable of infecting tobacco plants (Supplementary Figure 4).

The proposed structural destabilization mechanism offers the possibility of a cooperative disassembly reaction within the virion: the removal of a Ca^2+^ from its coordination site at lower radius has immediate effect on the neighboring Ca^2+^ sites, which are located in close proximity of 10 Å. Although the lower radius region of the virion is destabilized in low Ca^2+^ and basic environments, we find that the large part of the CP conformation is not affected by these environmental changes. This is an important aspect of the CP plasticity, which only requires a subtle destabilization of the metastable switch to trigger cotranslational disassembly [27] and, at the same time, to be sufficiently stable to initiate re-assembly of the virion for viral replication.

## Methods

### Sample preparation

TMV sample was isolated as described in [21] and stored in 0.1 M Tris-HCl pH 7.0, 0.02% NaN_3_ (w/v) at a concentration of ∼33 mg/ml at 4 °C. A total of 50 μl of virus stock solution was dialyzed for 1 hour at room temperature against 50 ml of 0.1 M NaOAc pH 5.2, 20 mM CaCl_2_ and 50 ml of MilliQ H_2_O, respectively. Before plunge-freezing sample concentration was adjusted to 22 mg/ml and 1.1 mg/ml for the Ca^2+^/acidic pH and the water condition, respectively. A total of 3.6 μl were applied on holey carbon grids (C-flat 300 mesh R2/2, Protochips) that had been glow discharged in an EasyGlow (Pelco) device. Grids were plunge-frozen in liquid ethane using a Vitrobot mark IV (Thermo Fisher Scientific) with a blotting time of 2 s at 10°C and 100% humidity.

### Electron Microscopy

Data acquisition was performed on a Titan Krios microscope (Thermo Fisher Scientific) operated at 300 kV, through a Gatan Quantum 967 LS energy filter using a 20 eV slit width in zero-loss mode. The dataset was recorded on a Gatan K2-Summit direct electron detector operated in super-resolution mode, at a calibrated magnification of 215,000 (resulting in a super-resolution pixel size of 0.325 Å on the object scale) with a defocus range of 0.15 – 0.35 μm. For the TMV in water, a total of 20 frames were recorded in movies of 5 s exposure at a dose rate of ∼2.6 e^−^/physical pix/s, accumulating a total dose of 30.8 e^−^/Å^2^ at the sample level. For the TMV in Ca^2+^/acidic pH conditions, a total of 40 frames were recorded in movies of 4 s exposure with a dose rate of ∼3.7 e^−^/physical pix/s, accumulating a total dose of 41.3 e^−^/Å^2^ at the sample level. For both samples, data collection was performed on a single grid using SerialEM [28].

### Image processing

After visual inspection of the micrographs, 62 images for the TMV in water, and 197 images for the TMV in Ca^2+^/acidic pH conditions, were selected and both datasets were processed in the same way. Briefly, movie frames were aligned and dose-compensated with MotionCor2 [29] using patch-based alignment (5 × 5) and Fourier space cropping (by a factor 2), resulting in a pixel size of 0.65 Å. Contrast transfer function parameters for the micrographs were estimated using Gctf [30]. Helix coordinates were determined automatically using MicHelixTrace [31], resulting in ∼20,000 segments for each sample. All 2D and 3D classifications and refinements were performed using RELION implementation of single-particle based helical reconstruction [6], including per-particle refinement of CTF parameters, correction of estimated beam tilt and “Bayesian polishing” [32]. Refined helical symmetry parameters were determined to a helical rise of 1.429 Å and a helical rotation of 22.036° for the TMV in water and 1.428Å/22.036° for the TMV in Ca^2+^/acidic pH conditions. The here determined helical rise of 1.429 Å (or 1.428Å) differs from previous studies, where 1.408 Å was used [5][21][2]. Reconstructions with starting values of a helical rise of 1.408 Å gave rise to typical artifacts from wrong symmetry and lack of high-resolution features [33]. To verify the obtained values, 3D maps were systematically evaluated using a series of pixels sizes after an additional round of unrestrained atomic model refinement. The resulting real-space cross correlation and EM ringer score were computed and the maximum was found for both criteria at a pixel size of 0.65 Å. This pixel size is in agreement with the results of the calibration procedure using Magical Reference Standard for TEM (Electron Microscopy Sciences). The reported overall resolutions for TMV of 2.1 Å in Ca^2+^/acidic pH and 2.0 Å in water conditions, were calculated using the Fourier shell correlation (FSC) 0.143 criterion. In parallel, we also processed the dataset taken in water with the single-particle based helical reconstruction package SPRING [33], which also gave rise to 2.0 Å resolution (Supplementary Figure 6). The final density maps were corrected for the modulation transfer function of the detector and sharpened by applying a negative B factor that was estimated using automated procedures [34] (−49 Å^2^ for the TMV in water and −68 Å^2^ for the TMV in Ca^2+^/acidic pH conditions). Local resolution maps were calculated with BlocRes [35] at a 0.5 FSC cutoff and the maps were subsequently locally filtered. To annotate significant molecular density in the 3D reconstruction and to control false positive voxels, confidence maps [25] were generated, where local resolution information was incorporated to increase statistical power, i.e. to decrease the amount of false negative voxels. Due to the large size of the reconstructed 3D volumes (450×450×450 voxels), quarters of the maps were cropped for further analysis and atomic modelling.

### Atomic model building and refinement

Atomic models were built and refined as 9-mers in order to account for inter-subunit interactions accurately. PDB *4udv* [5] was used as starting model and rigid body fitted into the processed maps using *Chimera* [36]. Additional H_2_O and Mg^2+^/Ca^2+^ ions were placed using the 1% FDR thresholded density. Several rounds of real-space refinement with *phenix.real_space_refine* [37] and manual rebuilding with *Coot* [38] were done to obtain the presented models. Refinement was performed using rotamer, Ramachandran and C-β restraints in addition to standard restraints of bond-lengths, angles, etc. Real-space refinement was carried out with global minimization together with a local grid search. Atomic coordinates and B-factors were refined against the sharpened and locally filtered maps. Residues 154-158 were not modelled as the corresponding density was not sufficiently well resolved to allow unambiguous atomic building. Additional significant density at 1% FDR at the N-terminus was modelled as a N-terminal acetylation, as reported in [39]. Grouped atomic displacement factors (ADP) were refined with *phenix.real_space_refine*. Validation scores were calculated with *phenix.molprobity* [40], *phenix.em_ringer* [41] and DipCheck [42]. To assess overfitting of the refinement, we introduced random coordinate shifts into the final models using the program *phenix.pdbtools*, followed by refinement against the first unfiltered half-map (half-map 1) with the same parameters as above. Comparisons of FSC curves of the randomized model refined against half-map1 versus half-map 1 and the FSC curve of the same model map versus half-map 2 do not indicate overfitting (Supplementary Figure 1b). Simulated model maps were calculated with *LocScale* [43].

### Infection of *Nicotiana tabacum* with TMV

The Bel B and Bel W3 variants of *Nicotiana tabacum* were chosen for infection experiments as they are known to be sensitive to TMV. Seeds distributed on a soil surface were watered and placed in a greenhouse for germination and cultivation at the following growth conditions: length of day: 14h, day: 28° C/ night: 22° C, relative humidity: 70%. Seedlings were piqued after 16 days, repotted after 31 days for the first time and repotted after 58 days for the second time. Before infection, the plants reached a height of 65 – 85 cm. On Day 60, one plant of both variants was infected with the TMV whereas the second plant was cultivated free of virus as a control. They were grown under ambient room temperature conditions and 16 hours of neon light. For infection, 25 µl of tobacco mosaic virus stock (33 mg/ml) was diluted into 10 ml PBS pH 7.5 and mixed with 105 mg Silicon carbide (SiC, 200-450 mesh, Sigma-Aldrich) in a porcelain mortar. SiC was used as an abrasive to cause small wounds and lesions supporting virus entry [44] [45]. The pestle was dipped into the virus/SiC suspension and rubbed gently onto the top surface of each plant leaf [46][47]. Ten days after the infection event, first symptoms of TMV replication were visible [48]. The variety Bel B developed deformed leaves, yellowish spots and new leaves were unusually light green and showed stunted growth. The variety Bel W3 showed lesions, necrotic spots on the leaves and stunted growth. After 35 days of infection, the plants possessed heights of 100 cm and 124 cm of Bel B and of Bel W3, respectively whereas the corresponding non-infected control plants were of 168cm and 167cm heights.

### Figure preparation

FSC and ADP graphics were visualized with ggplot2 in R [31][32]. Chimera [36] was used for the figure preparation of the molecular densities and atomic models and for preparation of the movie.

## Accession numbers

The EMDB accession numbers for the two cryo-EM maps and corresponding atomic coordinate models are EMD-xxx/EMD-yyy and PDB-xxx/PDB-yyy, respectively.

## Author contributions

FW, MB and CS designed research. FW prepared cryo-samples, acquired data and computed 3D reconstructions. MB built and interpreted atomic models. IvdH performed infection experiments. MB, FW and CS wrote the manuscript with input from IvdH.

## Acknowledgements

We are grateful to Thomas Hoffmann and Jurij Pecar (IT Services) for maintenance of the high-performance computing at EMBL. We thank Wim Hagen and Arjen Jakobi for discussions on the microscope magnification calibration. We would like to thank the Institute of Bio- and Geosciences - Plant Sciences (IBG-2) within the Forschungszentrum Jülich GmbH, especially Beate Uhlig, for providing expertise, material and space in the greenhouses to grow the tobacco plants.

## Competing financial interests

The authors declare that no competing financial interests exist.

---

**Supplementary Figure 1.**
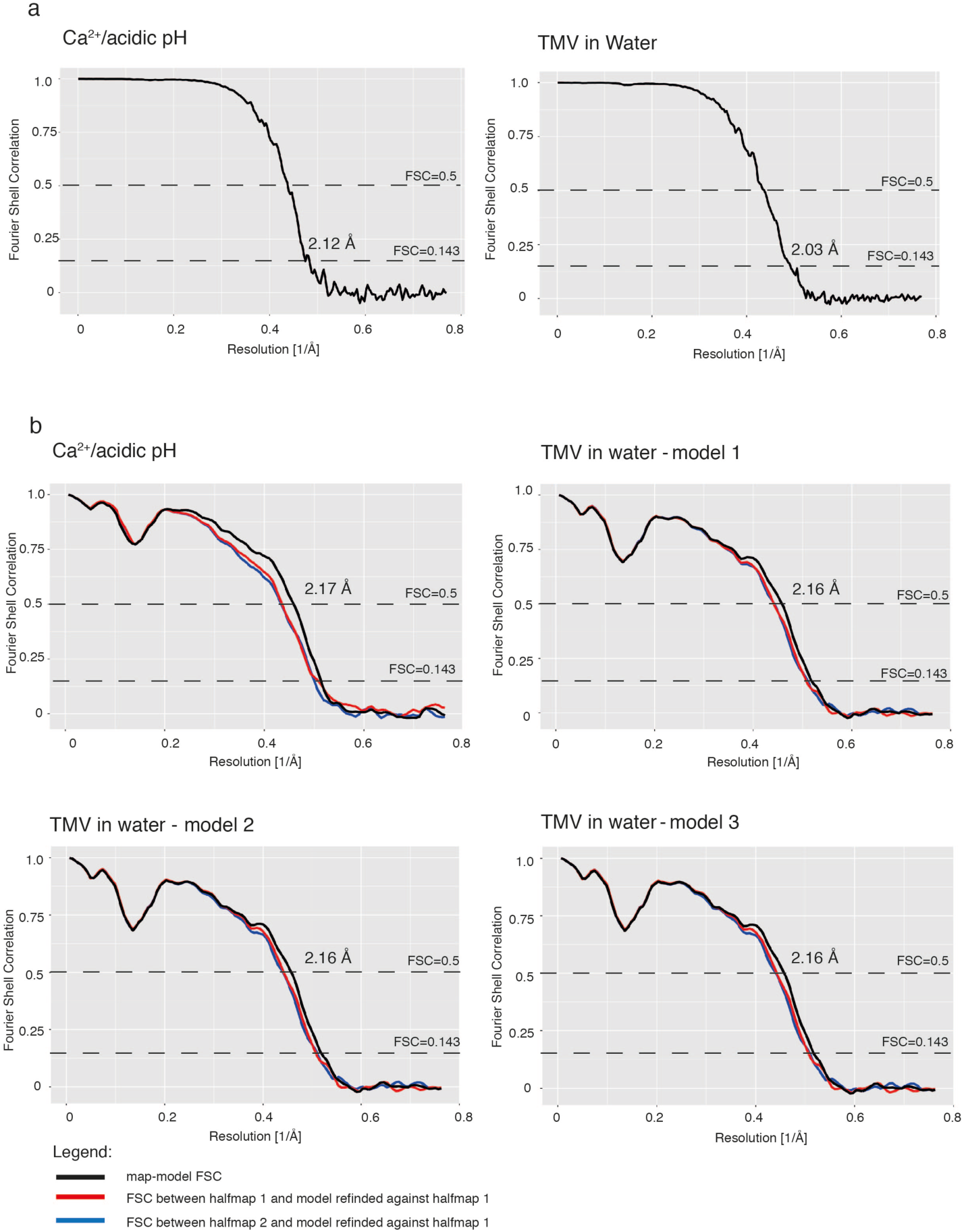
Resolution assessment using Fourier shell correlation (FSC). (a) Comparison of FSC curves between two half-maps for the Ca^2+^/acidic pH (left) and water structure (right). (b) FSC curves between map and model (black), between half map 1 and a perturbed model refined against half map 1 (red) as well as between half map 2 and a perturbed model refined against half map 1 (blue) for the 4 determined atomic models.

**Supplementary Figure 2.**
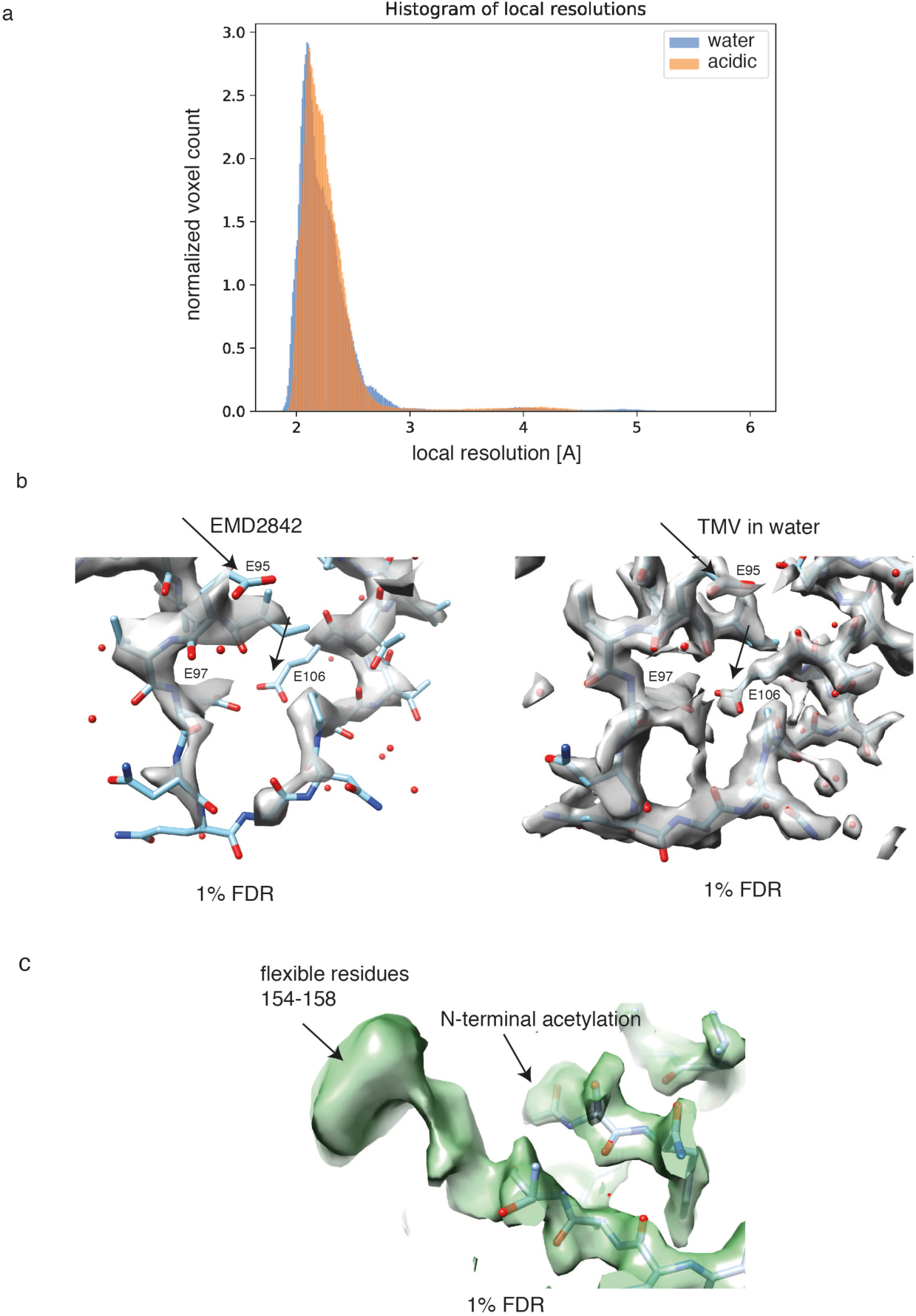
Local resolution assessment of Ca^2+^/acidic pH and water structure. (a) Overlay of local resolution histograms computed with BlocRes in the Ca^2+^/acidic pH (orange) and water condition (blue). Resolution of the water map is slightly higher. (b) Map comparison from previous study (left) [5] with this study in water (right) including overlaid current atomic model thresholded at a FDR of 1%. The here determined structure shows additional significant and defined density for the lower radius region. (c) Additional density for the flexible C-terminus and N-terminal acetylation are significant at an FDR of 1%.

**Supplementary Figure 3.**
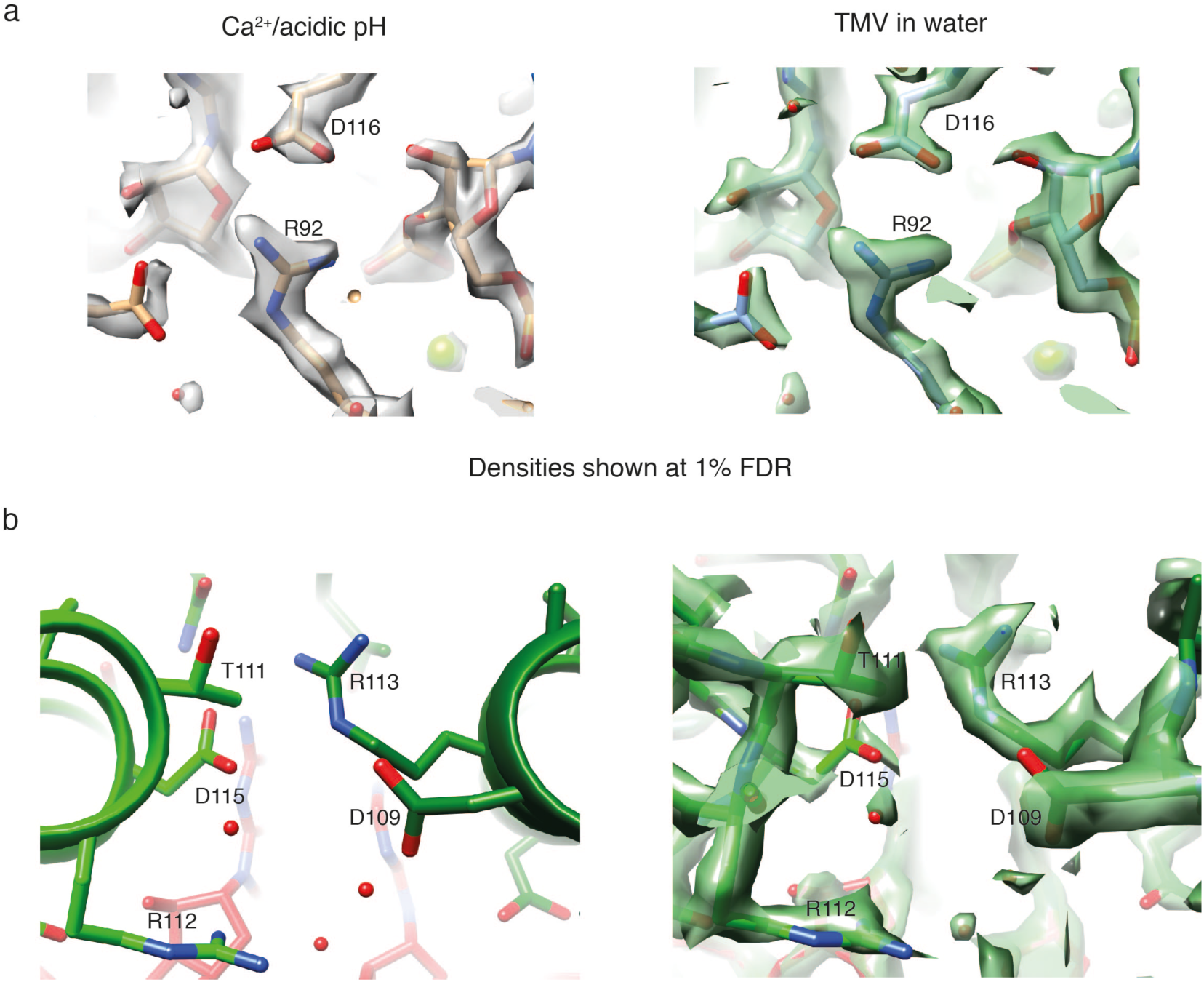
Structural details of Ca^2+^/acidic pH and water states. (a) Atomic models shown with the respective densities of TMV in Ca^2+^/acidic pH (left) and water condition (right) at the proposed location of a second Ca^2+^ site in proximity to the RNA [16][20]. No compatible Ca^2+^ ion density could be detected. (b) Residue D109 and its environment: no obvious interaction with other carboxylates is evident in our structures (left). The same view is shown with corresponding density at 1% FDR (right).

**Supplementary Figure 4.**
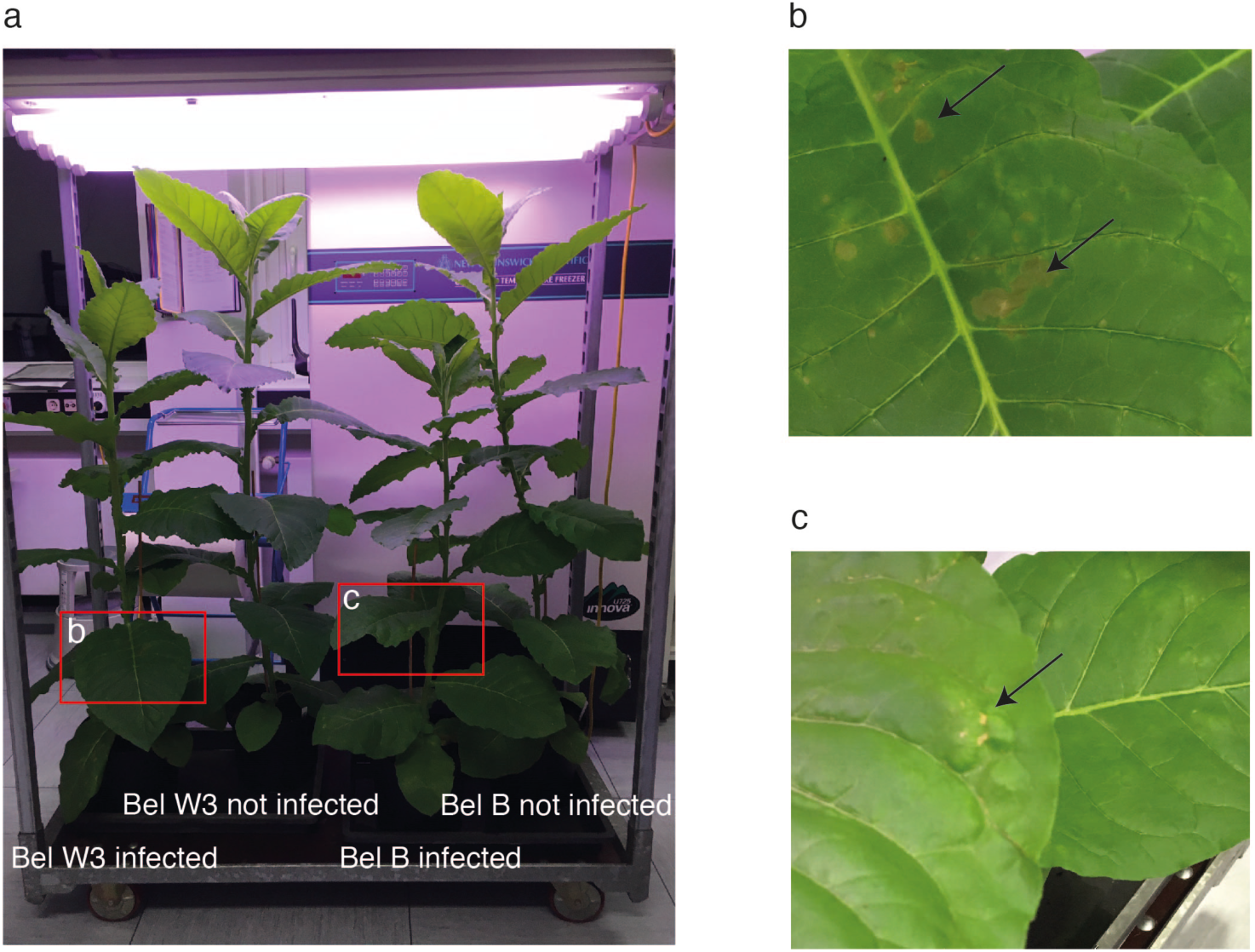
Symptoms of TMV infection on tobacco plants. (a) Four tobacco plants from left to right: variant Bel W3 infected, Bel W3 not infected, variant Bel B infected, Bel B not infected. Infected plants are significantly reduced in height in comparison with the non-infected control plants. (b) Leaf of variant Bel W3 with necrotic lesions (arrows). (c) Leaf of variant Bel B with light green spots (arrow) and bulges.

**Supplementary Figure 5.**
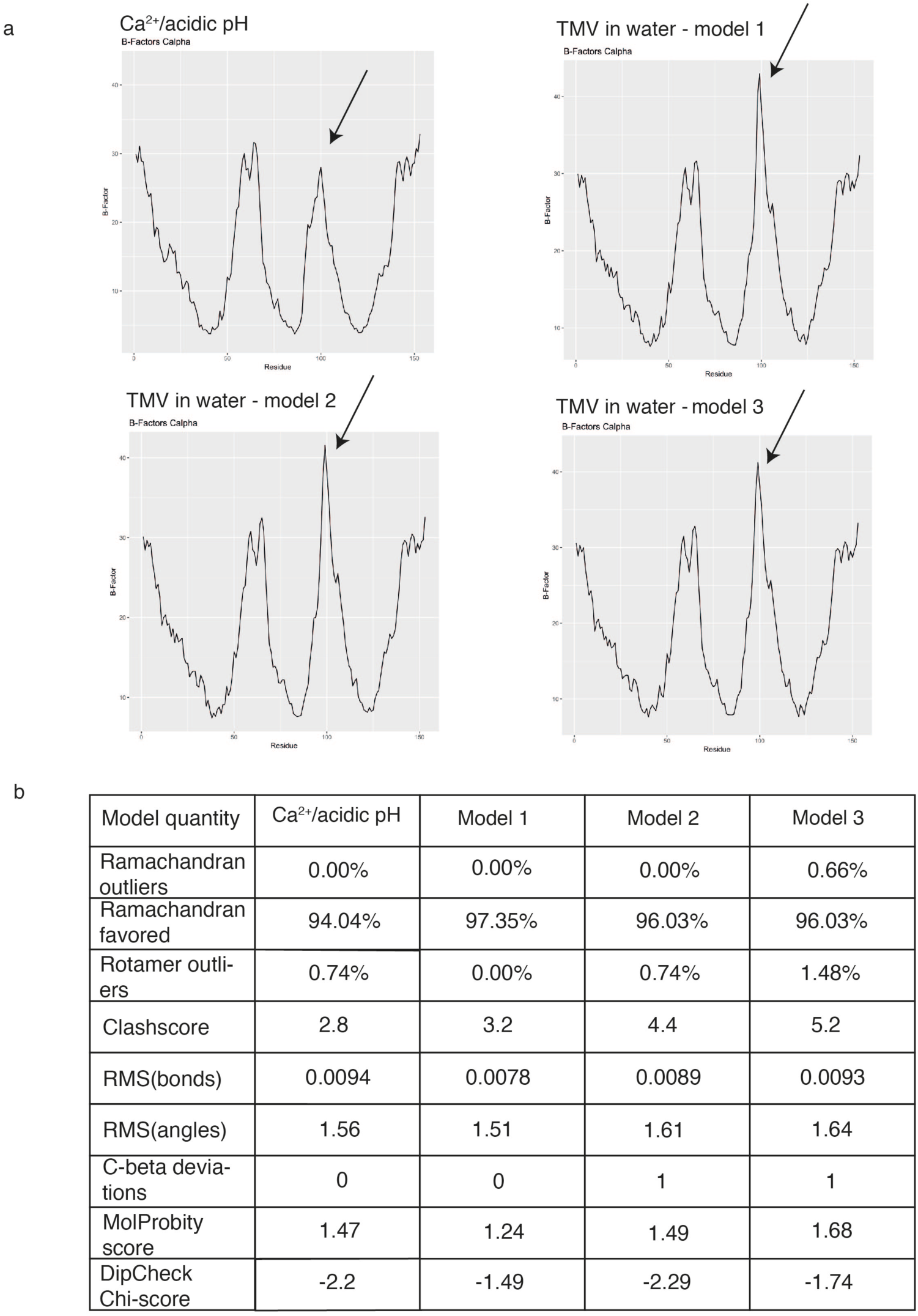
Analysis of refined atomic models. (a) Plot of Cα B-factors with corresponding residue number. Plot of 4 determined models from this study are shown. The peak for residues 90 to 110 at the lower radius region (highlighted with an arrow) shows lower B-factors in the Ca^2+^/acidic pH condition. For other residues, the overall profile is very similar. (b) Model validation statistics for all 4 refined models.

**Supplementary Figure 6.**
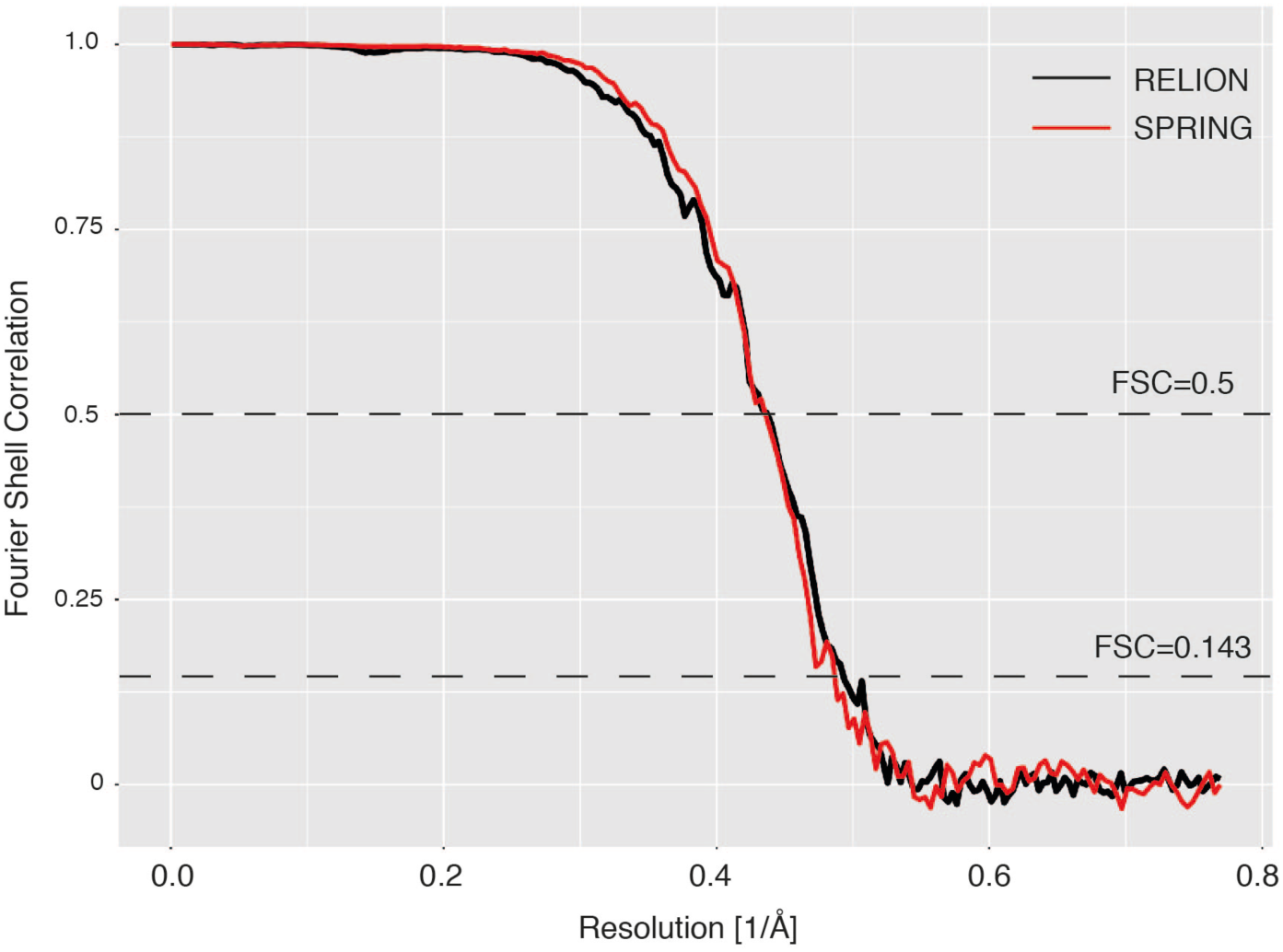
FSC curve comparison of RELION and SPRING reconstructions. Comparison of masked FSC curves of reconstructions determined with RELION (black) and SPRING (red) for TMV in water. Curves are in close overlap with each other at resolutions of 2.04 achieved with SPRING and 2.01 with RELION, respectively.

**Supplementary Movie 1. Structural transition of the determined Ca^2+^/acidic pH and water state.**

Movie shows Ca^2+^/acidic pH structure of the helical TMV assembly and subsequently zooms in the Ca^2+^ site with all residues involved in the switch mechanism highlighted in atom display. The structural transition was interpolated between the Ca^2+^/acidic pH model and model 1 of the water state. Model 1 showed the most different conformation compared with the Ca^2+^/acidic pH structure. Subsequently, all 3 models of the water structure are shown.

